# Deciphering Features of Metalloprotease Cleavage Targets Using Protein Structure Prediction

**DOI:** 10.1101/2025.01.13.632885

**Authors:** Dae Sun Chung, Jongkeun Park, Won Jong Choi, Dongwan Hong

## Abstract

**Introduction:** ADAM10 (A Disintegrin and Metalloproteinase 10) cleaves specific substrates, influencing diverse physiological and pathological processes. However, its substrate specificity and cleavage sites remain insufficiently characterized. This study aimed to identify and classify substrate features and elucidate cleavage sites using computational approaches.

**Methods:** Protein structure prediction was performed on 13 substrates with experimentally defined cleavage sites to analyze ADAM10–substrate interactions and assess their structural and spatial characteristics.

**Results:** Based on protein structure prediction of experimentally validated substrates, we identified four recurrent structural features associated with ADAM10-mediated cleavage. First, most substrates were predicted to interact with the proteolytically active form of ADAM10 (92.3%). Second, interaction sites were typically located in the extracellular region rather than intracellular or transmembrane domains (76.9%). Third, cleavage sites predominantly resided in unstructured or extended loop regions, corresponding to linear secondary structures (∼70%). Lastly, cleavage sites were spatially enriched in octants (1, 4, 5, and 8) relative to the catalytic Zn²□ ion in the active site (84.0%). These percentages were calculated per determinant. On the basis of these determinants, we developed a classification algorithm and organized 51 substrates accordingly. As a result, 82.4% of the substrates were assigned to either Group 1 (51.0%) or Group 2 (31.4%).

**Discussion:** This study presents a structure-informed classification approach for predicting ADAM10 cleavage substrates, enabling candidate identification without direct experimental validation.

**Conclusions:** We present a novel structure-based framework for classifying the substrates of ADAM10 and demonstrate its applicability to other Metalloproteases.

## 1 INTRODUCTION

Metalloproteases, including A Disintegrin And Metalloprotease (ADAM), are protease enzymes that contain a catalytic metal ion at their active site. ADAMs play a crucial role in regulating immune functions, particularly during inflammatory processes **[1, 2]**. Additionally, ADAMs have been implicated to various diseases, including cancer metastasis, inflammatory diseases, and neurological disorders **[3–6]**. The ADAM family is classified based on proteolytic activity into proteolytic ADAMs, non-proteolytic ADAMs, and pseudogenes. Proteolytic ADAMs include ADAM8, 9, 10, 12, 15, 17, 19, 20, 21, 28, 30, and 33, while non-proteolytic ADAMs include ADAM2, 7, 11, 18, 22, 23, 29, and 32 **[7]**.

ADAM family proteinases regulate the function of various cell surface proteins. ADAM10 cleaves substrates, such as Notch, amyloid precursor protein (APP), N-cadherin, E-cadherin, and Tumor-associated calcium signal transducer 2 (TACSTD2); all of which play critical roles in cancer cell proliferation, migration, and invasion **[8–14]**.

These properties suggest that ADAM10 serves as a promising target for cancer therapy via two distinct approaches. The first involves direct ADAM10 antigen targeting to inhibit cancer cell growth, potentially enhancing the efficacy of existing treatments in combination therapies **[15]**. The second focuses on targeting ADAM10 substrates, specifically the cleavage sites generated during proteolytic processes, which are associated with cancer cell growth, thereby suppressing tumor progression **[9–14]**.

These antigens hold considerable potential for use in antibody-drug conjugates (ADCs), which have recently gained attention. ADCs are a class of anticancer agents designed to deliver cytotoxic payloads directly to tumor sites by linking the payload to antibodies that specifically target tumor-associated antigens. This targeted approach results in reduced systemic toxicity compared to conventional chemotherapies **[16]**. However, antigen selection remains a major challenge in ADC development **[17]**. Given the highly potent cytotoxic agents utilized in ADCs, inappropriate antigen selection can lead to unintended off-target effects and an increased risk of drug resistance, which often hinder ADC development **[17, 18]**. Consequently, antigen selection for ADCs requires a nuanced, multifaceted approach that differs from traditional strategies **[19]**.

In the case of TACSTD2, a well-known example in ADC development, we employed a multifaceted approach that utilizes the cleavage sites of ADAM10 substrates, such as TACSTD2, which is critical for cancer cell growth, as antigens. Although ADAM10, a proteolytic enzyme, has great potential as a novel cancer therapeutic target, the number of candidate substrates cleaved by ADAM10 remains limited **[8]**.

For cleaved substrates, cleavage sites have been identified based on diverse characteristics. For example, in furin, a specific consensus motif within the substrate protein sequence mediates interaction between furin and its substrates **[20]**. Tucher et al. also identified the cleavage sites of ADAM10 and ADAM17 through mass spectrometry-based profiling. However, their analysis revealed no detectable sequence similarity among different substrates **[21]**.

Consequently, we explored alternative sequence-based and amino acid property-based features, but none were sufficient for reliable identification under these constraints. This led us to adopt a structure-informed strategy, leveraging scalable AI-predicted models to capture the native context of protease substrate interactions.

In this study, we therefore aimed to explore the medical applications that have emerged following recent advancements in AI-based protein structure prediction technologies, such as AlphaFold and RosettaFold. **[22, 23]**. These software tools have yielded promising results in predicting protein structures **[24]**. Given their primary application in protein binding, we recognized the potential of these technologies to analyze cleavage processes, specifically the interaction between a protease and its substrate. To address these challenges, we developed a prediction algorithm focused on the proteins cleaved by ADAM10, utilizing the structure predictions generated by these tools.

Overall, we aimed to predict protein structure by utilizing the structural characteristics of ADAM10, classifying candidate substrates based on cleavage-related features, and developing an algorithm to facilitate this process.

## 2 MATERIALS AND METHODS

### 2.1 Data collection

To obtain the structural and sequence information for ADAM10, we accessed data from UniProt (https://www.uniprot.org/) using the ADAM10 entry (O14672). Additionally, we identified the sequence and positional information for the various ADAM10 domains, including the Signal peptide (S, 1-19), Pro-domain (Pro, 20-213), Metalloprotease domain (MP, 214-456), Disintegrin domain (Dis, 457-551), Cysteine-rich domain (Cys, 552-672), Transmembrane domain (M, 673-693), and Cytoplasmic domain (Cyto, 694-748) **(Figure 1A) [25]**. Similarly, we identified the sequence and positional information of the Nitrogen epsilon 2 (NE2) atoms of HIS383, HIS 387, and HIS393, which are amino acid residues involved in metal ion binding, which is associated with cleavage **[26]**.

**FIGURE 1.**
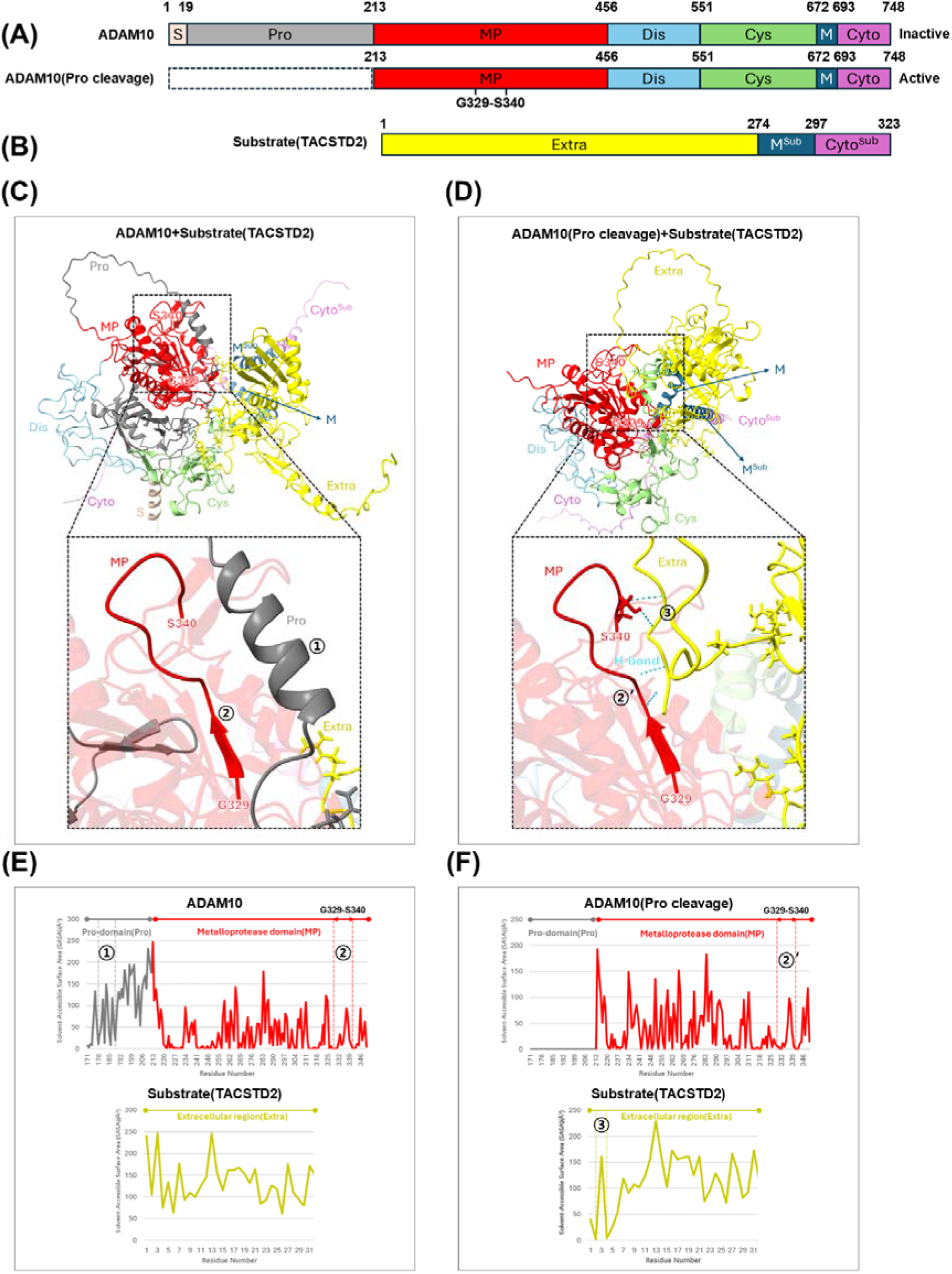
Identification of ADAM10 composition, substrate structure, and common binding region of ADAM10. **(A):** Domain composition of ADAM10. S: Signal sequence(Light orange), Pro: Pro-domain(Gray), MP: Metalloprotease domain(Red), Dis: Disintegrin domain(Sky blue), Cys: Cystein-rich domain(Light green), M: Transmembrane region(Navy blue), Cyto: Cytoplasmic region(Purple), ADAM10(Pro cleavage) represents the protease-active state of ADAM10, where the Pro-domain, which inhibits the protease function, has been removed. The G329-S340 region in the MP domain of ADAM10(Pro cut) is the sequence that commonly binds to substrates exclusively in the Pro cleavage state. **(B):** Domain composition of the representative ADAM10 substrate (TACSTD2). Extra: Extracellular region(Yellow), MSub: Transmembrane region of the substrate(Navy blue), CytoSub: Cytoplasmic region of the substrate(Purple). **(C):** Predicted structure of the ADAM10+Substrate (TACSTD2) complex. The enlarged section shows that, due to the presence of the Pro-domain (17), no interaction occurs between the G329-S340 region (17) of ADAM10 and the substrate (TACSTD2). **(D):** Predicted structure of the ADAM10(Pro cleavage)+Substrate (TACSTD2) complex. The enlarged section shows that the G329-S340 region (17’) of ADAM10(Pro cleavage) interacts with the substrate (TACSTD2) (17), forming hydrogen bonds (H-bonds). **(E):** SASA graph of the enlarged region of the ADAM10+Substrate (TACSTD2) complex. The vertical axis represents SASA values, and the horizontal axis represents amino acid residue numbers. The mutually low SASA values of the Pro-domain (17) and the G329-S340 region (17) of ADAM10 indicate interactions between these regions, which, through the high SASA values of the substrate (TACSTD2), confirm that interaction with region 17 is prevented. **(F):** SASA graph of the enlarged region of the ADAM10(Pro cleavage)+Substrate (TACSTD2) complex. The low SASA values of the G329-S340 region (17’) of ADAM10(Pro cleavage) and the substrate (TACSTD2) (17) confirm their interaction.

We obtained the sequence and cleavage site information for proteins cleaved by ADAM10, not only for TACSTD2, but also by using data provided by Biochem et al. (2009) **[27]**. These substrates include Amyloid-beta precursor protein (APP), Probetacellulin (BTC), Cadherin-1 (CDH1), Cadherin-2 (CDH2), Delta-like protein 1 (DLL1), Pro-epidermal growth factor (EGF), Tumor necrosis factor ligand superfamily member 6 (FASLG), Low affinity immunoglobulin epsilon Fc receptor (FCER2), Proheparin-binding EGF-like growth factor (HBEGF), Interleukin-6 receptor subunit alpha (IL6R), Neurogenic locus notch homolog protein 1 (NOTCH1), and Melanocyte protein PMEL (PMEL). In addition, 38 other substrates were included in the literature review. The exact cleavage sites of the added substrates are unknown **(Table 1)**. The sequences and positional information of the substrates were identified using UniProt, using a similar process as followed for ADAM10.

**Table 1.**
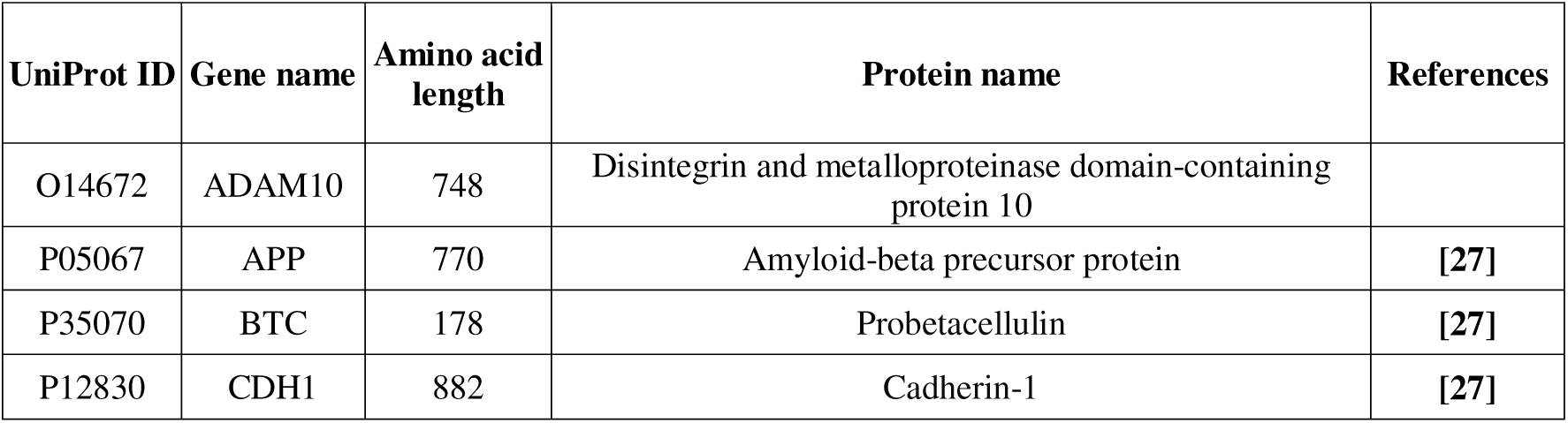
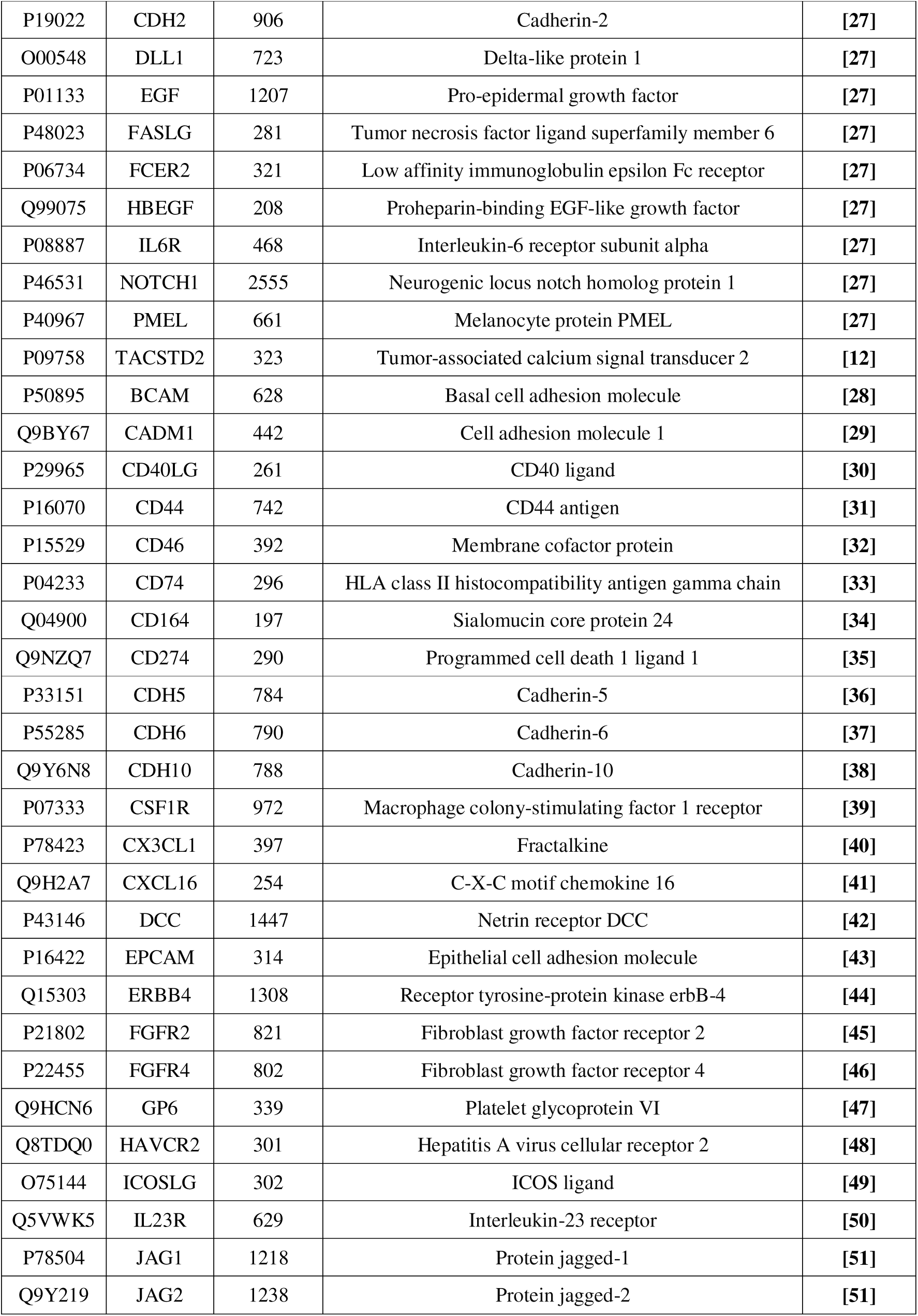
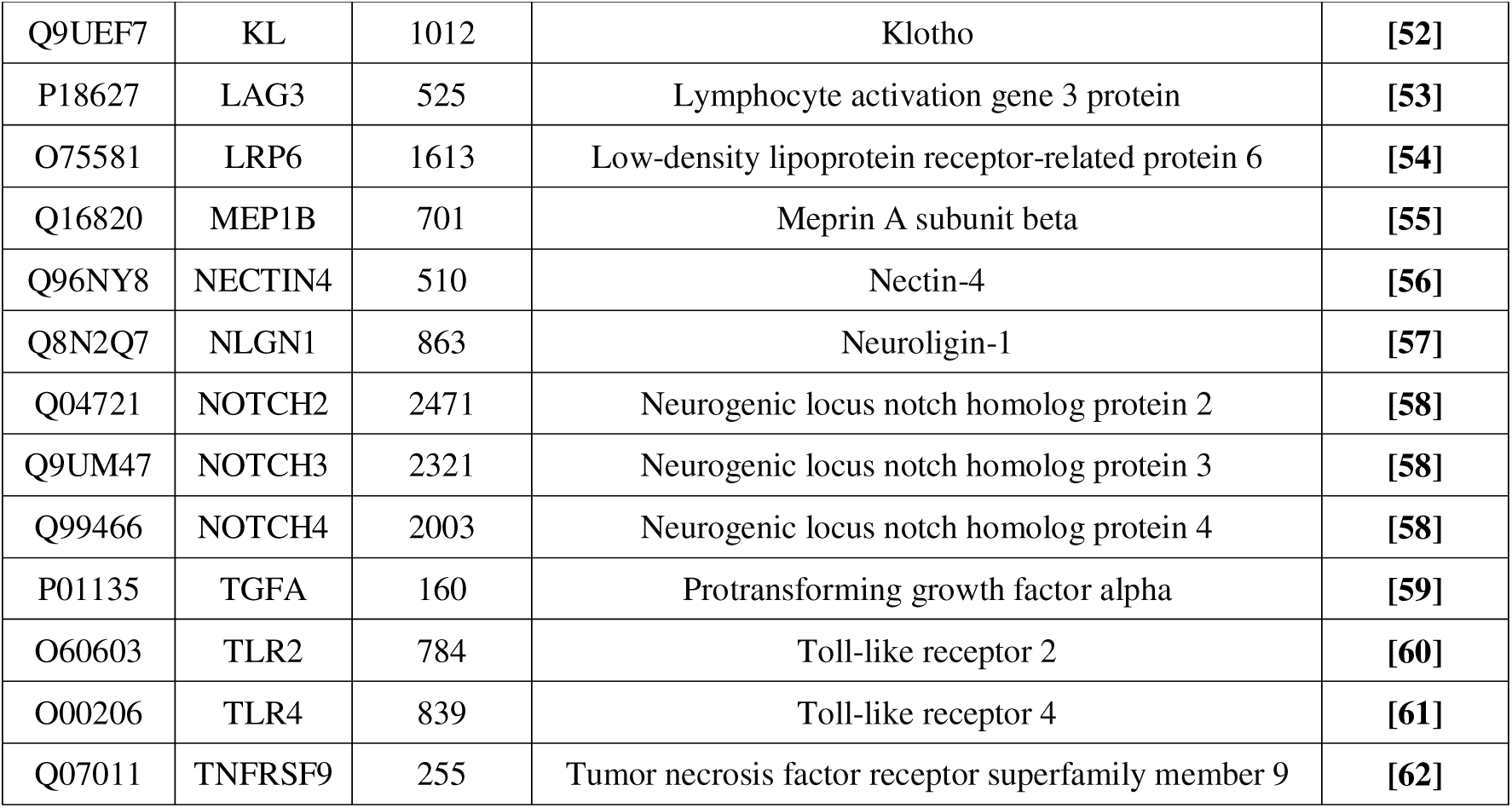
Information on ADAM10 & ADAM10 substrates.

### 2.2 Protein structure prediction

To determine the interactions between ADAM10 and its substrates after cleavage, we examined both the precursor form, in which the Pro-domain sequence remains intact, and the mature form, in which the Pro-domain is removed **(Figure 1A)**. The sequence of the Pro-domain is as follows:

“N-QYGNPLNKYIRHYEGLSYNVDSLHQKHQRAKRAVSHEDQFLRLDFHAHGRHFNLRMKRDTSLFSDEFKVETSNKVLDYDTSHIYTGHIYGEEGSFSHGSVIDGRFEGFIQTRGGTFYVEPAERYIKDRTLPFHSVIYHEDDINYPHKYGPQGGCADHSVFERMRKYQMTGVEEVTQIPQEEHAANGPELLRKKR -C”.

To analyze the structure of proteins with and without the Pro-domain, we utilized a system running Ubuntu 22.04.4 LTS with Linux kernel version 6.5.0-44-generic and used AlphaFold2 (AF2) release 2.3.2 for structural predictions. For the prediction of multimeric structures between ADAM10 and candidate substrates, we set the "model_preset=multimer" flag. Next, to improve the prediction quality, we used random seeds. High-confidence regions were consistently predicted, regardless of the random seed, while low-confidence regions were not. Therefore, we explored various structural possibilities by adjusting the initial conditions using random seeds. This approach allowed for the retrieval of Protein Data Bank (PDB) files consisting of five ranked models, with one PDB file per rank by setting the "num_multimer_predictions_per_model=1" flag. Finally, to further enhance the stability and quality of the predicted structures, we applied the Assisted Model Building with the Energy Refinement (AMBER) relaxation process to all PDB models by setting the "models_to_relax=all" flag **(Figure S1)[63]**

To verify the protein structures, Chimera X (version 1.6.1) was used to explore the spatial structures obtained from the PDB files generated by AF2.

### 2.3 Analysis of multimeric protein (ADAM10, Substrate) PDB files

The predicted PDB files, generated based on the identified sequence information, contain essential data, such as chains identifiers distinguishing the two proteins, atom names, and atomic coordinates. These files were analyzed further using various computational algorithms to gain deeper insights. First, hydrogen bond interactions between ADAM10 and its substrate, as well as intra-substrate hydrogen bonds, were identified using Biopython software. Subsequently, the secondary structures of the proteins (α-helix, β-sheet) and Solvent-Accessible Surface Area (SASA) were analyzed using the DSSP algorithm **[64]**. SASA is a robust tool for quantifying and describing Protein-Protein interactions (PPI). A SASA value approaching zero indicated that the corresponding protein region was involved in these interactions **[65]**.

To evaluate the predicted structures, we used the UniProt identifiers for each substrate to retrieve experimentally validated structures from the Protein Data Bank (www.rcsb.org). The root-mean-square deviation (RMSD) was then calculated to compare the experimental and predicted structures **[66]**. RMSD is a widely used metric for assessing the similarity between two protein structures, with lower values indicating greater similarity. An RMSD value below 2 Å suggests a high degree of structural similarity, making it an ideal measure for evaluating the accuracy of predicted protein structures **[67]**.

All the analyses were performed using Python (version 3.11.9), Biopython (version 1.83), or mkDSSP (version 4.0.4).

## 3 RESULTS

### 3.1 Identification of ADAM10 composition, substrate structure, and common binding region of ADAM10

We investigated the domain structure of ADAM10, and the structural changes induced upon cleavage. In its protease-inactive form, ADAM10 consisted of seven domains: S, Pro, MP, Dis, Cys, M, and Cyto. Upon cleavage of the Pro-domain, the protease-active form of ADAM10 consisted of MP, Dis, Cys, M, and Cyto **(Figure 1A)**. TACSTD2, an ADAM10 substrate, was composed of an extracellular region (Extra), a transmembrane region (M^Sub^), and a cytoplasmic region (Cyto^Sub^), which were also present in other substrates **(Figure 1B)**.

Based on these findings, we used AlphaFold-Multimer to predict and analyze the complex structures of ADAM10 and its 13 substrates **[12, 27]**, comparing their protease activity-dependent interactions. In the presence of the Pro domain, no interactions were observed between the GLY329-SER340 region of ADAM10 and TACSTD2, owing to steric hindrance caused by the Pro domain. However, upon Pro-domain removal, hydrogen bond interactions were observed between ALA331, VAL333, SER339, and SER340 of ADAM10 and ALA2, GLY4, and PRO5 of TACSTD2 **(Figure 1C and 1D)**.

Upon examining the remaining substrates, we observed consistent interactions in the GLY329–SER340 region of ADAM10 for 11 substrates, excluding CDH2, in the absence of the Pro domain **(Figure S2-S3, Table S1)**.

To further confirm these findings quantitatively, we measured the SASA. The results indicated that, in the presence of the Pro-domain, the SASA values for the Pro-domain region (designated as region ①) were VAL178 (11 Å²), MET182 (14 Å²), and GLN186 (8 Å²). For the GLY329-SER340 region of ADAM10 (designated as region ②), the SASA values were ALA331 (0 Å²), VAL333 (34 Å²), SER339 (22 Å²), and SER340 (10 Å²) **(Figure 1E)**. In contrast, in the absence of the Pro-domain, the SASA values for the GLY329-SER340 region (designated as region ②’) were ALA331 (0 Å²), VAL333 (12 Å²), SER339 (31 Å²), and SER340 (4 Å²), while the SASA values for TACSTD2 (designated as region ③) were ALA2 (1 Å²), GLY4 (4 Å²), and PRO5 (25 Å²) **(Figure 1F)**.

Similarly, SASA measurements of the remaining 11 substrates, excluding CDH2, showed comparable patterns in their interactions with ADAM10 **(Table S2-S5)**. These results quantitatively demonstrated that these substrates shared the characteristic of binding to the GLY329-SER340 region of protease-active ADAM10.

Next, we assessed the feasibility of actual binding between GLY329-SER340 and various substrates. This region resided within the 1–672 domain **(Figure 1A)**, which is external to the membrane, indicating that for substrate binding to occur, the binding site must also be located in the Extra.

In TACSTD2, the Extra spanned residues 1–274 **(Figure 1B)**, with binding sites for ALA2, GLY4, and PRO5 located within this Extra. Therefore, we confirmed that binding between the GLY329-SER340 region and TACSTD2 was feasible. Similarly, among the 11 substrates that bind to the GLY329-SER340 region, nine substrates, excluding NOTCH1 and DLL1, were confirmed to bind within the Extra, further supporting the feasibility of these interactions **(Figure S3, Table S6-S7)**.

### 3.2 Structural features of cleavage sites in substrates

We examined the structures of the cleavage sites to predict and characterize the substrate features. The ARG87-THR88 site of TACSTD2 was cleaved by ADAM10 **(Figure 2A) [12]**. To explore this, we conducted a structural investigation of TACSTD2. The SER81-VAL90 region containing the cleavage site was observed to have a linear structure **(Figure 2B)**.

**FIGURE 2.**
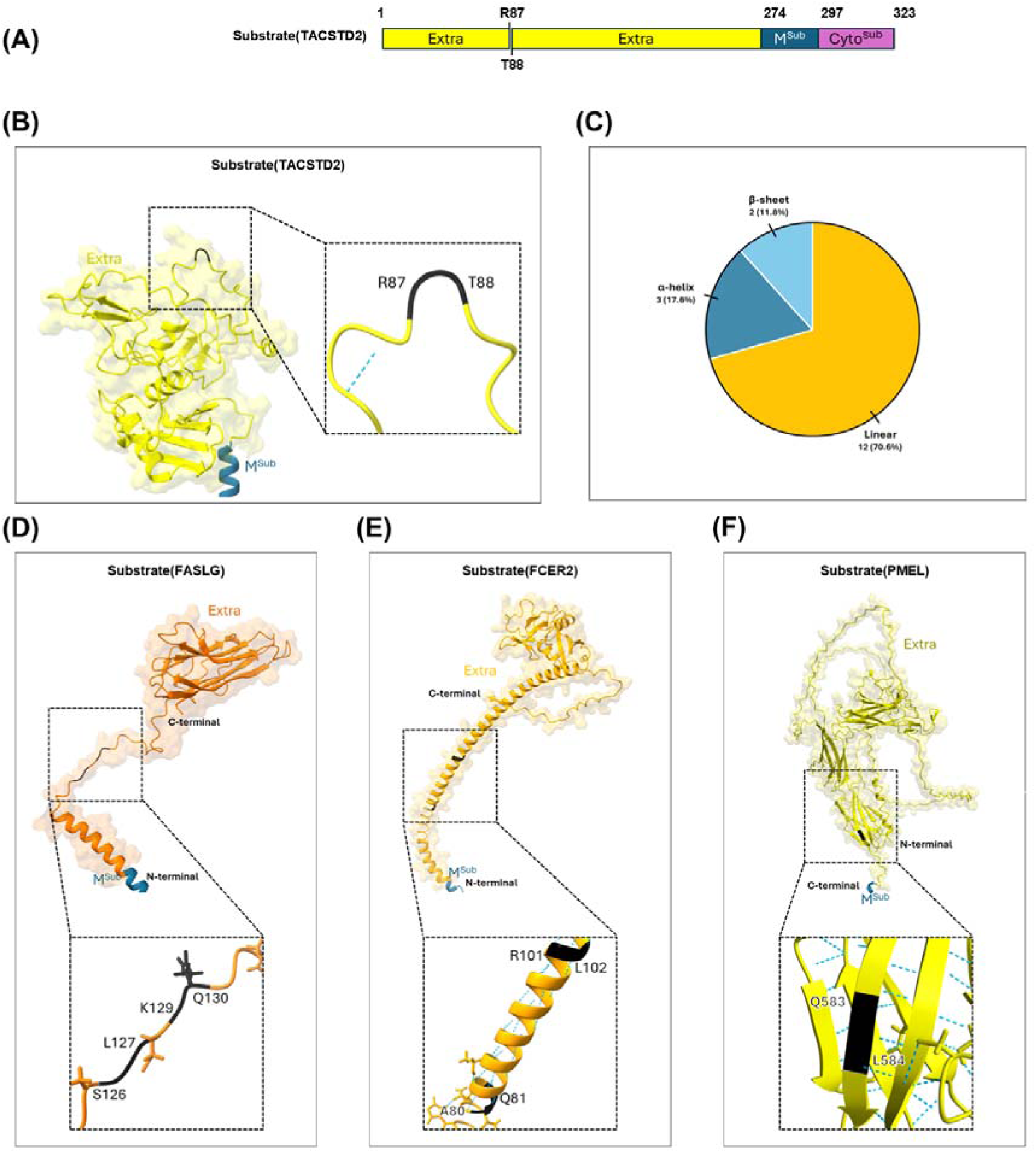
Structural Features of Cleavage Sites in ADAM10 Substrates. **(A):** Domain composition of the representative ADAM10 substrate (TACSTD2), including its cleavage site. **(B):** Predicted structure of the cleavage sites R87 and T88 of the substrate (TACSTD2) cleaved by ADAM10. The cleavage site of the substrate (TACSTD2) was predicted to adopt a linear structure. **(C):** Pie chart of the structural characteristics of substrate cleavage sites. Among the 17 cleavage sites analyzed, 12 were predicted to adopt a linear structure, 3 were α-helix structures, and 2 were β-sheet structures. **(D):** Predicted structure of the cleavage sites S126 and L127, and K129 and Q130 of the substrate (FASLG). The cleavage sites of the substrate (FASLG) were predicted to adopt a linear structure. **(E):** Predicted structure of the cleavage sites A80 and Q81, and R101 and L102 of the substrate (FCER2). The cleavage sites of the substrate (FCER2) were predicted to adopt an α-helix structure. **(F):** Predicted structure of the cleavage site Q583 and L584 of the substrate (PMEL). The cleavage site of the substrate (PMEL) was predicted to adopt a β-sheet structure.

We verified the cleavage sites of the remaining 12 substrates and found that BTC, EGF, FASLG, and FCER2 each possessed two cleavage sites **(Table 2) [27]**.

**Table 2.** Cleavage sites targeted by ADAM10.

| Gene | Number(P1) | Number(P1') | P5 | P4 | P3 | P2 | P1 |  | P1' | P2' | P3' | P4' | P5' |
| --- | --- | --- | --- | --- | --- | --- | --- | --- | --- | --- | --- | --- | --- |
| APP | 687 | 688 | V | H | H | Q | K | - | L | V | F | F | A |
| BTC | 42 | 44 | S | P | E | T | N | (G) | L | L | C | G | D |
|  | 111 | 112 | V | D | L | F | Y | - | L | R | G | D | R |
| CDH1 | 700 | 701 | R | K | A | Q | P | - | V | E | A | G | L |
| CDH2 | 714 | 715 | T | D | V | D | R | - | I | V | G | A | G |
| DLL1 | 535 | 536 | D | L | T | E | K | - | L | E | G | Q | G |
| EGF | 970 | 971 | H | Y | S | V | R | - | N | S | D | S | E |
|  | 1023 | 1024 | W | W | E | L | R | - | H | A | G | H | G |
| FASLG | 126 | 127 | H | T | A | S | S | - | L | E | K | Q | I |
|  | 129 | 130 | S | S | L | E | K | - | Q | I | G | H | P |
| FCER2 | 80 | 81 | G | D | Q | M | A | - | Q | K | S | Q | S |
|  | 101 | 102 | A | E | Q | Q | R | - | L | K | S | Q | D |
| HBEGF | 149 | 150 | G | L | S | L | P | - | V | E | N | R | L |
| IL6R | 357 | 358 | S | L | P | V | Q | - | D | S | S | S | V |
| NOTCH1 | 1720 | 1721 | Y | K | I | E | A | - | V | Q | S | E | T |
| PMEL | 583 | 584 | V | V | S | T | Q | - | L | I | M | P | G |
| TACSTD2 | 87 | 88 | P | K | N | A | R | - | T | L | V | R | P |

In total, we examined the structures of 17 cleavage sites identified across all 13 substrates, including TACSTD2 **(Figure 2C)**. Twelve of these cleavage sites, including TACSTD2 (ARG87-THR88), FASLG (SER126-LEU127), and FASLG (LYS129-GLN130), exhibited linear structures **(Figure 2B, D)**. Three cleavage sites, including FCER2 (ARG80-GLN81) and FCER2 (ARG101-LEU102), were observed to adopt an α-helix structure **(Figure 2E)**. Finally, two cleavage sites, including PMEL (GLN583-LEU584), were observed to adopt a β-sheet structure **(Figure 2F) (Figure S4-S5)**. Over 70% of cleavage sites exhibited linear structures **(Figure 2C)**.

The predicted structures were further evaluated using RMSD **(Table S8)**. Notably, CDH2 was excluded from analysis due to the absence of experimentally validated structural data.

For TACSTD2, the RMSD between the experimentally determined structure, the crystal structure of the extracellular part of human TACSTD2 (PDB ID: 7PEE), and the predicted structure was 0.774 Å. With an RMSD value lower than 2 Å, this indicates a high degree of structural similarity. Similarly, the remaining 11 substrates also exhibited RMSD values below 2 Å for at least one experimentally determined structure, demonstrating a strong structural alignment with the predicted models **[66, 68]**.

We then examined whether the experimentally determined cleavage site structures matched the predicted cleavage site regions. The 7PEE structure revealed that the actual cleavage site exhibited a linear structure, which is consistent with our predictions **[68]**.

Similar validations were performed for other substrates. Notably, only APP and NOTCH1 had experimental data available for their cleavage sites. APP exhibited both the predicted linear structure and a secondary structure at the cleavage site, while NOTCH1 showed a β-sheet structure, which are also consistent with our predictions. For the remaining substrates, no experimental data were available for the cleavage sites or findings unrelated to the cleavage sites **(Table S8)**.

Subsequently, we investigated whether regions with fewer hydrogen bonds among neighboring amino acids were more likely to be cleavage sites by analyzing the number of hydrogen bonds in and around each cleavage sites. For the cleavage site in TACSTD2, ARG87 formed one hydrogen bond, while THR88 formed none. Furthermore, an examination of hydrogen bonds in the adjacent ALA82-THR96 and LEU89-VAL90 regions revealed the following: ALA82, LEU89, and VAL90 each had zero hydrogen bonds; PRO83, LYS84, and ASN85 each had one, and THR86 had two hydrogen bonds. For the remaining 12 cleavage sites with linear structures (excluding the five cleavage sites with secondary structures, such as α-helix or β-sheet, that generally require multiple interactions with surrounding residues), we also counted hydrogen bonds in the surrounding amino acids **(Table S9)**. We found that the cleavage sites in linear structures typically had two or fewer hydrogen bonds with their surrounding amino acids.

### 3.3 Spatial characteristics between cleavage sites and the zinc ion

The extracellular domain of ADAM10 consisted of MP, Dis, and Cys domains, while PRO, M, and Cyto domains were excluded from the full structure. Within the MP domain, key histidine residues (HIS383, HIS387, and HIS393) were involved in binding zinc ions, which play critical roles in proteolytic cleavage **(Figure 3A) [69]**.

**FIGURE 3.**
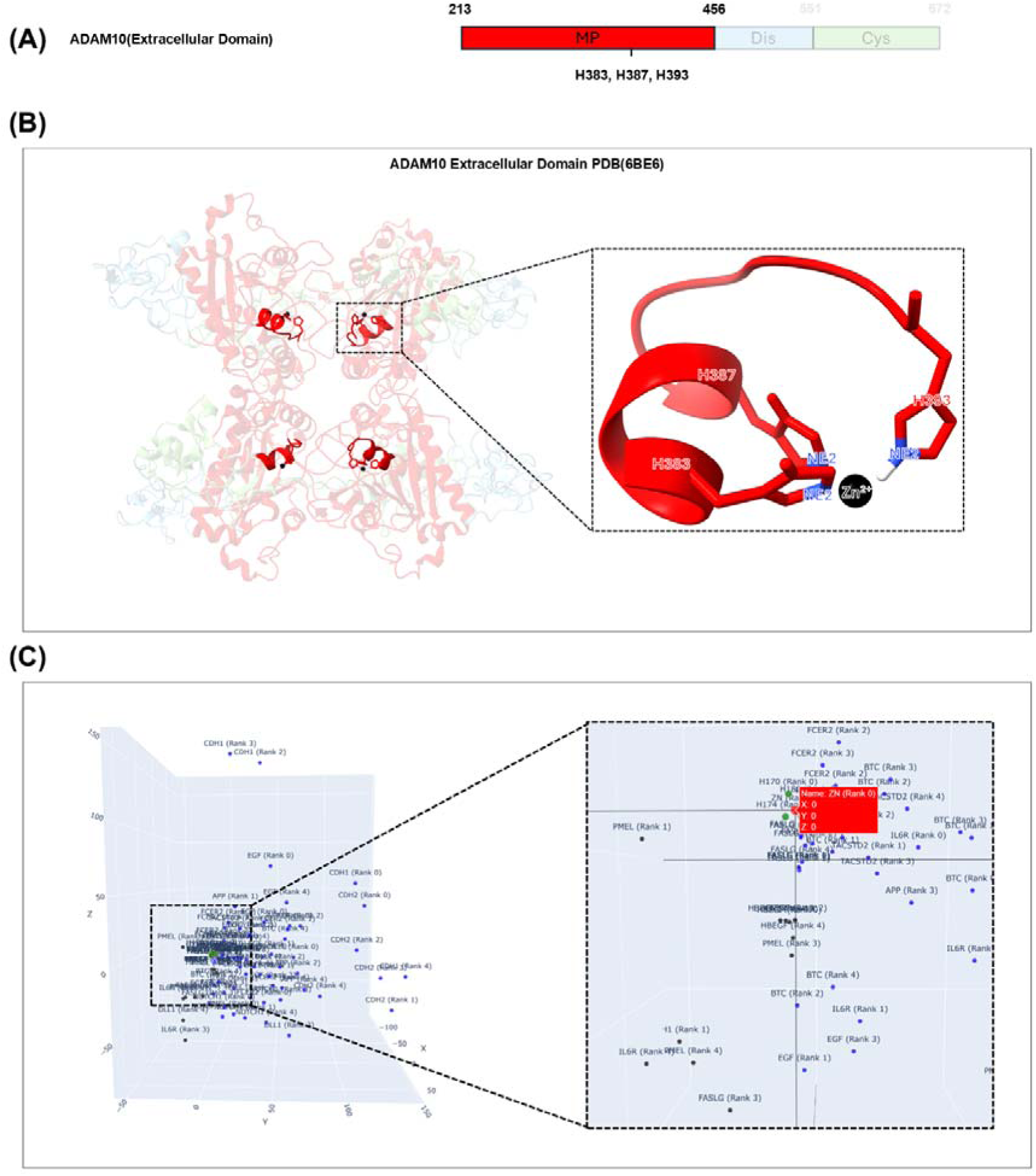
Spatial Characteristics between Cleavage Sites and the Zinc ion. **(A):** Metalloprotease domain (MP) in the extracellular domain of ADAM10. The MP contains three histidines (H383, H387, H393) that serve as binding sites for the zinc ion, a metal ion directly involved in proteolytic cleavage. **(B):** The PDB structure of ADAM10 Extracellular Domain (PDB ID: 6BE6), showing the spatial relationship between ADAM10 and the zinc ion. The enlarged section highlights the zinc ion and the Nitrogen epsilon 2 (NE2) atoms of histidines H383, H387, and H393 that coordinate its binding. **(C):** Coordinate distribution of 85 cleavage sites (17 cleavage sites × five ranked) aligned with the zinc ion as the origin (0, 0, 0). Among these, 71 cleavage sites were found to be located within Octants 1, 4, 5, and 8, excluding one site from CDH1, one from DLL1, two from FASLG, five from HBEGF, two from IL6R, and three from PMEL.

Our aim was to investigate the spatial characteristics of the cleavage sites in the substrates, focusing on the zinc ions associated with the proteolytic activity of ADAM10. First, we predicted the location of the zinc ion in the PDB file of ADAM10. We used the PDB structure of the ADAM10 Extracellular Domain (PDB ID: 6BE6) **(Figure 3B)**. The 6BE6 structure consisted of four ADAM10 extracellular domains, each containing a zinc ion. We measured the distances between the zinc ion and NE2 atoms of the three histidines involved in its coordination: HIS383, HIS387, and HIS393. The distances between the zinc ion and HIS383 NE2 were 2.134 Å, 2.189 Å, 1.998 Å, and 2.106 Å. For HIS387 NE2, the distances were 2.058 Å, 2.082 Å, 2.082 Å, and 2.062 Å. The distances for HIS393 NE2 were 2.058 Å, 2.100 Å, 2.341 Å, and 2.290 Å.

Based on the measured distances, we predicted the location of the zinc ions in the previously predicted PDB structures of the substrates **(Figure 3B)**. To explore the spatial relationship between the zinc ion and the cleavage sites, we visualized the coordinates of the zinc ion along with those of 85 cleavage sites (derived from 17 cleavage sites across the five ranked models). The coordinates were aligned such that the zinc ion was positioned at the origin (0,0,0) **(Figure 3C)**.

The cleavage sites of TACSTD2 were located at the following coordinates: (−18.38, 20.39, 19.67), (−19.83, 10.43, −4.92), (−35.42, 9.41, 1.39), (−32.55, 12.96, −7.15), and (−25.38, 17.17, 3.34). The cleavage sites of the other substrates were positioned at similar coordinates (Figure S5, Table S10).

The results revealed that, with the exception of one site in CDH1, one in DLL1, two in FASLG, five in HBEGF, two in IL6R, and three in PMEL, a total of 71 cleavage sites fell within the coordinate range of −72.83 to 49.01, 0.02 to 145.59, −55.91 to 161.78. This indicates that over 84% of the cleavage sites were located in octants 1, 4, 5, and 8 relative to the zinc ion at the origin **(Figure S5-S6)**.

### 3.4 Classification of substrates and development of an algorithm based on identified features

Using the aforementioned characteristics, we classified the substrates into two categories: 13 substrates with known cleavage sites (represented in black) and 38 substrates with unknown cleavage sites (represented in red) as input data **(Figure 4A)**.

**FIGURE 4.**
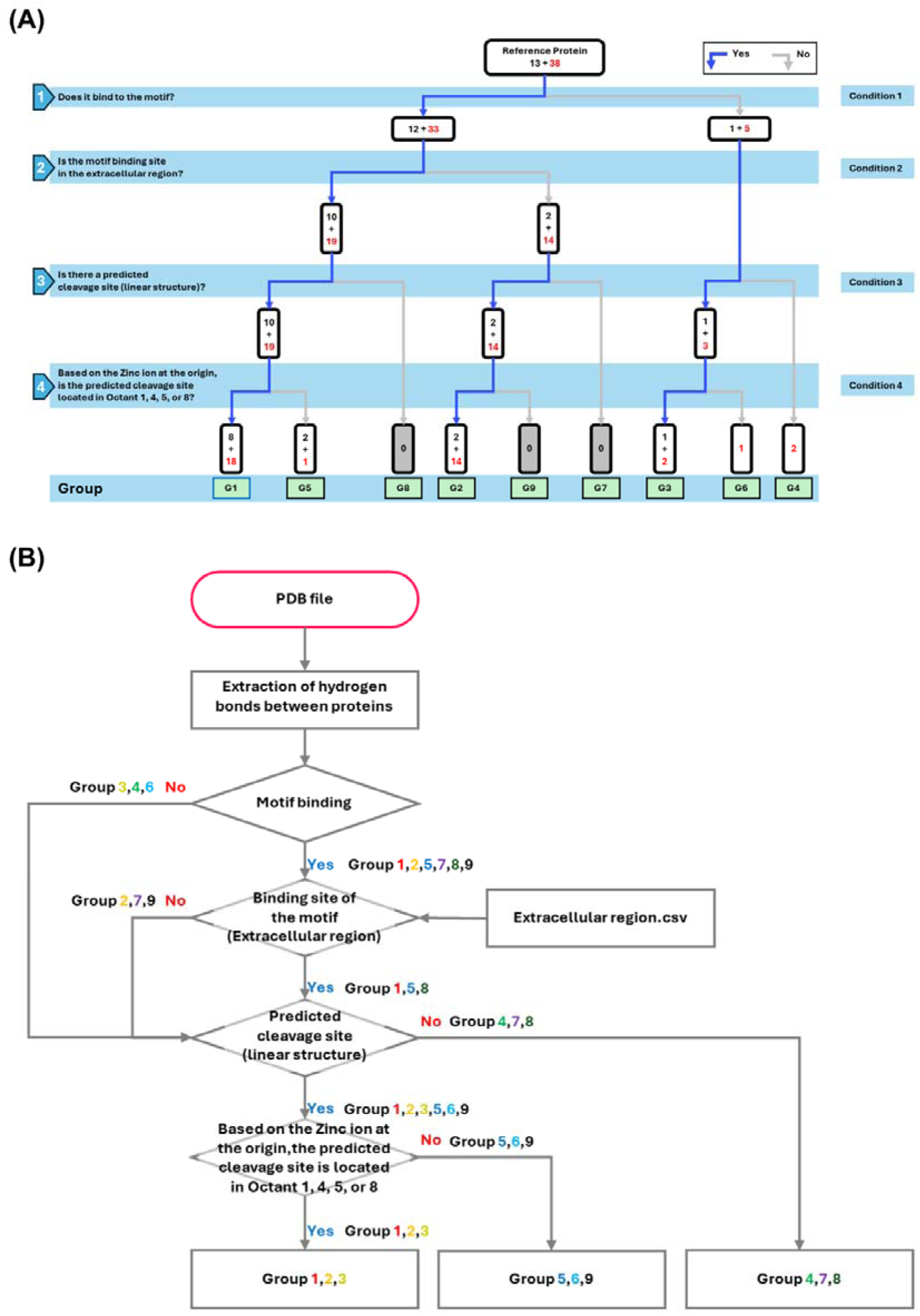
Classification of Substrates and Algorithm Development Based on Identified Features. **(A):** Decision tree showing the classification of ADAM10 substrates based on specific conditions. The classification was performed on a total of 51 substrates, including 13 substrates with known cleavage sites (represented in black) and 38 substrates for which cleavage sites remain unknown (represented in red). **(B):** Flowchart depicting the classification process of the substrates.

First, we used protein structure prediction to model the complex structures of ADAM10 and its substrates, generating PDB files for further analysis. Next, we extracted hydrogen bond interactions between ADAM10 and its substrates from the PDB files to determine whether the substrates bound to a common binding region of ADAM10. If no binding was detected, the substrates were classified into Groups 3, 4, and 6. In contrast, if binding was observed, the substrates were classified into Groups 1, 2, 4, 5, 7, 8, and 9. For the binding groups, we further evaluated whether the substrates were bound to the Extra.

If binding occurred in the Extra, the substrates were classified into groups 1, 5, and 8; and if binding did not occur in the Extra, the substrates were classified into groups 2, 7, and 9. Next, we assessed whether the substrates contained predicted cleavage sites with the structural features similar to known cleavage sites based on hydrogen bonding and structural data within the substrates. If no predicted cleavage sites were identified, the substrates were classified into Groups 4, 7, or 8. To determine the spatial orientation of the predicted cleavage sites relative to the zinc ion, we evaluated whether the cleavage sites in Groups 1, 2, 3, 5, 6, and 9 were positioned within octants 1, 4, 5, and 8. If this condition was met, the substrates were classified into Groups 1, 2, or 3; otherwise, they were classified into Groups 5, 6, or 9 **(Figure 4B)**.

Among the 51 substrates analyzed, hydrogen bond formation with the common binding region of ADAM10 was observed in 45 substrates, whereas 6 showed no such interaction **(Table S1 and Table S11)**. 30 substrates were confirmed to bind within the extracellular region **(Table S6–S7 and Table S12–S13)**. In addition, 49 substrates harbored either experimentally validated or computationally predicted cleavage sites **(Table S14)**, with 45 of these positioned within octants 1, 4, 5, and 8 relative to the catalytic zinc ion **(Table S10 and Table S15)**.

Consequently, 26 of the 51 substrates (51%; which satisfied all identified features) were classified into Group 1. While the binding region of the substrates in Group 2 was predicted to fall outside the feasible binding region within the common binding site in ADAM10, 16 substrates were classified into Group 2 (31.4%; these substrates met all other criteria). The remaining substrates were classified into their respective Groups based on the respective criteria **(Figure 4A) (Table S16)**.

## 4 DISCUSSION

In the present study, we aimed to predict potential antigen cleavage candidates using ADAM10 (a disintegrin and metalloproteinase). While the role of ADAM10 in tumors and its cleavage effects are well established, information the identification of antigens cleaved by ADAM10 remains limited **[70]**. In addition, mechanisms underlying membrane protein localization on the cell membrane remain unknown, and experimental validation of these cleavage events is challenging due to technical limitations. Therefore, in this study, we propose a novel substrate-classification approach to predict cleavage candidates. To operationalize this approach, substrates were partitioned into nine groups using four structural conditions **(Figure 4)**: Group 1 (Conditions 1–4), Group 2 (Conditions 1, 3, 4), Group 3 (Conditions 3, 4), Group 4 (none), Group 5 (Conditions 1–3), Group 6 (Condition 3), Group 7 (Condition 1), Group 8 (Conditions 1, 2), and Group 9 (Conditions 1, 3).

Using this classification approach, we found that substrates with known cleavage sites (8) and those with unknown cleavage sites (18) were classified as Group 1. Additionally, known cleavage sites (2) and unknown cleavage sites (14) were classified as Group 2, indicating that approximately 82.4% of the substrates were categorized into either Group 1 or Group 2.

The primary difference between these two groups is that Group 2 does not fulfill Condition 2 (extracellular-binding condition).

Additionally, for this group, examination of the 16 substrates revealed that, apart from CD164, CSF1R, FGFR4, and IL23R, the remaining 12 are involved in cell–cell interactions, mediating direct intercellular contacts **(Table S17)[71–78]**.

These findings indicate that substrates involved in cell-cell interactions were more likely to be classified as Group 2 rather than Group 1.

Membrane-associated antigens play a critical role in tumor progression. For instance, ERBB2 (HER2) is expressed in breast and gastric cancer and serves as an important therapeutic target **[79]**. Similarly, CD79b has become an important target in cancer therapies **[80]**. Recent advances in ADCs have focused on targeting these antigens. Notably, trastuzumab deruxtecan, an ADC targeting ERBB2 (HER2), and polatuzumab vedotin-piiq, an ADC targeting CD79b, have been FDA approved **[81]**. TACSTD2, another membrane protein cleaved by ADAM10, has been identified in its cleaved form in tumors **[12]**, and ADCs targeting these cleavage sites are currently undergoing clinical trials, and have shown promising results **[82]**.

We considered how proteases recognize and cleave their substrates. A well-known example is furin, which recognizes and cleaves substrates at a short consensus sequence (R-X-[K/R]-R↓). In the case of metalloproteases such as ADAM10 and ADAM17, which are the focus of our study, cleavage site selection also appears to involve consensus-like motifs. Quantitative cleavage profiling revealed a preference at the P1’ position: ADAM10 tends to cleave substrates with leucine or aromatic residues, whereas ADAM17 shows a preference for valine. However, these findings suggest that consensus sequences alone are insufficient to reliably identify cleavage sites. Therefore, a structural understanding of the substrate is essential for accurate site prediction. To capture the structural features and patterns around cleavage sites, AI-based protein structure prediction methods may offer a powerful approach to model the complex architectures of substrates.

To identify the antigens cleaved by metalloproteases, we used a multistep approach: First, we utilized protein structure prediction techniques to extract PPI and identify secondary structural features (such as linear regions, α-helices, and β-sheets) **[21]**. Second, we analyzed the three-dimensional spatial distribution of cleavage sites based on physical distances in 3D space **[83]**.

Through this approach, our study provides a novel perspective, demonstrating that cleavage event predictions can be achieved not only through experimental methods **[12, 27]**, as in previous studies, but also via computational approaches. Furthermore, the recent release of AlphaFold 3 and similar advancements suggest that such methods will continue to improve, offering even greater potential for future research **(Figure S7) [84]**.

Nevertheless, our study has a few limitations. First, limited data were available for training and validation. The limited number of predicted protease-substrate complex structures and known substrates may have introduced potential biases. Moreover, the exact cleavage sites were not explicitly defined in the experimental data, which hindered our availability to identify consensus sequences of amino acids at these cleavage sites **(Figure S4 and S8)**. This is because our study did not seek to define a sequence-level consensus motif but instead examined the surrounding structural context in which cleavage is likely to occur.

Consequently, cleavage site predictions are provided as regions rather than specific positions. Additionally, RMSD analysis of the predicted structures revealed that the similarities with NMR and EM structures were lower than those with X-ray structures. This result highlights a limitation in the dataset, as 92% of the experimentally validated structural data in the PDB were derived from X-ray crystallography, resulting in insufficient NMR and EM data available for training AlphaFold 2 models **[85]**.

Second, evaluating analytical performance with conventional metrics is not feasible in our setting. Such metrics require well-defined True Positive (TP), True Negative (TN), False Positive (FP), and False Negative (FN) counts. However, our dataset consists of 51 literature-supported ADAM10 substrates and lacks a verifiable negative set because non-cleavage cannot be established experimentally across contexts. Accordingly, rather than focusing on statistical accuracy or recall, this study sought to classify the currently known cleaved substrates.

Third, limitations arise from biological complexity and cellular context. Due to the in silico nature of the analysis, cellular context such as extracellular and intracellular environments was excluded. Consequently, the interpretation relied exclusively on the predicted tertiary structures of the proteins.

## 5 CONCLUSION

In conclusion, phenomena that previously required exclusively experimental approaches ADAM10 substrate interactions and the ensuing cleavage can be addressed computationally: by leveraging AI-based protein structure prediction to predict putative cleavage regions and recapitulate the process in silico, we can effectively identify cleavage sites.

## LIST OF ABBREVIATIONS

ADAM10 A: Disintegrin and Metalloproteinase 10
AF2: AlphaFold2
SASA: Solvent-Accessible Surface Area
PDB: Protein Data Bank
RMSD: The Root Mean Square Deviation
TP: True Positive
TN: True Negative
FP: False Positive
FN: False Negative
ALA: Alanine (A)
GLY: Glycine (G)
ILE: Isoleucine (I)
LEU: Leucine (L)
CYS: Cysteine (C)
MET: Methionine (M)
PHE: Phenylalanine (F)
PRO: Proline (P)
THR: Threonine (T)\
SER: Serine (S)
TRP: Tryptophan (W)
TYR: Tyrosine (Y)
VAL: Valine (V)
HIS: Histidine (H)
ASN: Asparagine (N)
ASP: Aspartic acid (D)
GLU: Glutamic acid (E)
GLN: Glutamine (Q)
ARG: Arginine (R)
LYS: Lysine (K)

## ETHICS APPROVAL AND CONSENT TO PARTICIPATE

Not applicable.

## CONSENT FOR PUBLICATION

Not applicable.

## AVAILABILITY OF DATA AND MATERIALS

Not applicable.

## ACKNOWLEDGEMENTS

We thank the Global Science experimental Data hub Center (GSDC) and the Korea Research Environment Open NETwork (KREONET) service for data computing and network provided by the Korea Institute of Science and Technology Information (KISTI) and the HPC Innovation Hub of Telecommunications Technology Association (TTA). This work was supported by the ICT R&D program of MSIP [R7518-16-1001, Innovation hub for High Performance Computing].

## AUTHOR CONTRIBUTION

DC and JP designed the experiments, performed the experimental analysis, and wrote the original draft. DC, JP, and WC conducted the artificial intelligence analysis. WC contributed to data collection. DC, and JP were responsible for writing, review, and editing. DH supervised the study and assisted in drafting the manuscript. All authors read and approved the final manuscript.

